# Immune activation during broadly neutralizing antibody-mediated HIV suppression prior to post-intervention control

**DOI:** 10.64898/2026.05.29.728863

**Authors:** Julia A. Wagner, Demi A. Sandel, Ravi K. Patel, George W. Gruenhagen, Rafael Tibúrcio, Shayleen S. Singh, Kaiti Schwartz, Michela Traglia, Pamela Milani, Clara Di Germanio, Shlomi Ilan, Thomas Dalhuisen, Rebecca Hoh, Meghann C. Williams, Michiko Shimoda, Reuben Thomas, Boris Juelg, Matthew H. Spitzer, Amelia N. Deitchman, Michael P. Busch, Gabriela K. Fragiadakis, Peter W. Hunt, Michael J. Peluso, Steven G. Deeks, Rachel L. Rutishauser

## Abstract

Broadly neutralizing antibodies (bNAbs) have been associated with enhancement of HIV-specific T or B cell responses and sustained partial control of HIV replication in some people with HIV (PWH). The mechanisms through which bNAbs may potentiate host immunity in this context are not known. We previously reported the outcomes of a clinical trial in which ten PWH on antiretroviral therapy (ART) received a combination of immunotherapies including two bNAbs administered immediately preceding an analytic treatment interruption (ATI). After bNAb levels waned, seven participants exhibited varying degrees of post-intervention control of HIV linked to a robust expansion of activated CD8+ T cells in response to rebounding virus. To investigate the role of the bNAbs in enhancing endogenous immune responses, we looked for evidence of HIV-specific or broader immune activation during the period after ART was paused and prior to rebound when bNAbs were controlling HIV replication. At a timepoint early post-ART interruption and at least one month before virus emerged in plasma, we detected an increase in levels of plasma inflammatory proteins as well as phenotypic and transcriptional activation of innate and adaptive immune cells. Compared to non-controllers, post-intervention controllers demonstrated unique transcriptional activation patterns as well as differential longitudinal plasma inflammatory protein trends. No enhancement of HIV-specific T cell or antibody responses was observed in this window. This study identifies activated cell types and inflammatory pathways that are recruited early during bNAb-mediated HIV suppression and that may play a role in potentiating long-lasting HIV immune control after bNAb therapy.

**One Sentence Summary:** In a combination immunotherapy trial with high rates of post-intervention control, bNAb-mediated HIV suppression was associated with increased immune activation compared to HIV suppression by ART.

## INTRODUCTION

Monoclonal antibodies (mAbs) are used for the treatment of a variety of clinical conditions, including infections, malignancies, and autoimmune diseases (1). Beyond their well-established neutralizing and non-neutralizing functions, mAb:antigen immune complexes can potentiate long-lasting, antigen-specific T and B cell responses even after the antibodies wane, a phenomenon known as the “vaccinal effect” (2–6).

Broadly neutralizing antibodies (bNAbs) are mAbs that recognize the HIV envelope glycoprotein (Env) and confer cross-clade viral neutralization (7). Combinations of infused bNAbs have been used in HIV prevention, treatment, and cure (8). In several HIV cure studies involving bNAbs administered to people with HIV (PWH) either at the time of antiretroviral therapy (ART) initiation/re-start or at the time of pausing ART in a monitored analytic treatment interruption (ATI), some degree of post-ART HIV control was observed in up to 30-40% of participants after the bNAb levels waned (9–14).

The mechanisms that promote enhanced host immune responses after mAb infusion are well-established in animal models of acute viral infection and cancer. mAb:antigen immune complexes can directly interact with B cells and/or the constant (Fc) region of the mAbs can interact with innate immune cells expressing Fc receptors (FcR). In mice, these complexes can be taken up by antigen-presenting cells and presented to stimulate T and B cell responses, resulting in the generation of antigen-specific adaptive immunity (2,4,6,15). In non-human primates, bNAb administration shortly after simian HIV (SHIV) infection induces CD8+ T cell-mediated viral control in approximately 50% of animals (16), and CD8+ T cell-mediated SHIV control has been described when bNAbs/interleukin (IL)-15 were administered prior to an ATI (17). bNAb interaction with HIV virions may be important for potentiating host immune responses, as much lower rates of post-bNAb control have been observed when bNAbs are administered during suppressive ART (18).

The mechanisms by which bNAbs may potentiate host immune responses in humans are less well-defined. In PWH, bNAb administration has been shown to boost HIV Gag-specific CD8+ T cell magnitude and/or proliferative responses (9,10,12,14), though these increases were mostly small in magnitude, transient, and not correlated with HIV control, and do not always occur even when HIV control after bNAb administration is observed (19). bNAb administration has also been linked to autologous anti-HIV Env antibody induction (20,21) and to reservoir reduction (11,21). Therefore, there are several gaps in our understanding of how bNAbs administered to PWH may potentiate host immune responses to the virus to induce what is now termed “post-bNAb control.”

Our group previously reported the results of a combination immunotherapy study involving a T cell-based therapeutic vaccine as well as two infusions of two bNAbs (10-1074 (22) and VRC07-523LS (23)), with the second infusion given immediately preceding an ATI (13). Seven out of ten participants demonstrated enhanced control of HIV after the interventions with undetectable virus for greater than 18 months (1 participant) or low average viral loads (~1,000 copies/mL) sustained for several months (6 participants) after bNAb levels waned. Post-intervention control of HIV was linked to an increased expansion of activated and cycling CD8+ T cells in the blood in response to rebounding virus. While study participants showed evidence of broad innate and adaptive immune cell activation immediately prior to rebound, this activation occurred similarly in controllers and non-controllers. Therefore, the contribution of the bNAbs to the post-intervention control observed in this clinical trial remains unknown.

In this study, we hypothesized that we would observe evidence of immune activation during bNAb-mediated HIV suppression that could potentiate HIV-specific adaptive immune responses in participants in the combination immunotherapy trial. To this end, we applied a suite of sensitive cellular and plasma assays to first investigate whether we could detect evidence of adaptive and/or innate immune activation and enhancement early following ATI and prior to quantifiable rebound. We also investigated whether the immune activation we observed was related to low-level, residual viremia during this period. Finally, we asked whether the post-intervention control observed in this study was associated with differential immune activation during this critical window.

## RESULTS

### Study cohort and sample timepoints

A schema of the UCSF-amfAR HIV combination immunotherapy clinical trial is depicted in **Fig. 1A** and full clinical trial details have recently been published (13). Briefly, the Phase I single-arm study enrolled ten people with HIV on suppressive ART who were given a combination of immunotherapies including a therapeutic vaccine regimen during ART suppression and two bNAbs (10-1074 and VRC07-523LS) immediately preceding an ATI. Based on data from other cohorts, we expect that ART levels in participants in our study would have waned from the plasma by 10 days post-ATI (24). HIV rebound – defined as the first of two HIV-1 RNA measurements that were quantifiable on a clinical assay – occurred a median of 16.4 (range: 5.7-25.9) weeks after ART was paused. For the present study, we mostly focused on the following timepoints to characterize immune activation during the period of bNAb-mediated viral suppression: *preATI* (on ART, sampled within a week prior to ATI), *bNAb* (a median of 25 [range: 24-52] days post-ATI/bNAb infusion and ≥28 days before rebound), *preR* (pre-rebound, a median of 15 [range: 5-28] days prior to rebound in participants who experienced rebound), and *postR* (post-rebound, the first peripheral blood mononuclear cell [PBMC] samples available, collected ≤28 days after rebound and at a low viral load [<2,600 HIV-1 RNA copies/mL for PBMC samples; <6,500 copies/mL for plasma samples] in participants who experienced rebound). We previously reported that levels, cumulative exposure, and phenotypic susceptibility testing of the two bNAbs against rebound virus isolates correlated with time to rebound but not post-intervention control (13). See **Supp. Table 1** for participant-level demographic and clinical data and **Supp. Table 2** for viral load rebound curve data.

**Fig. 1:**
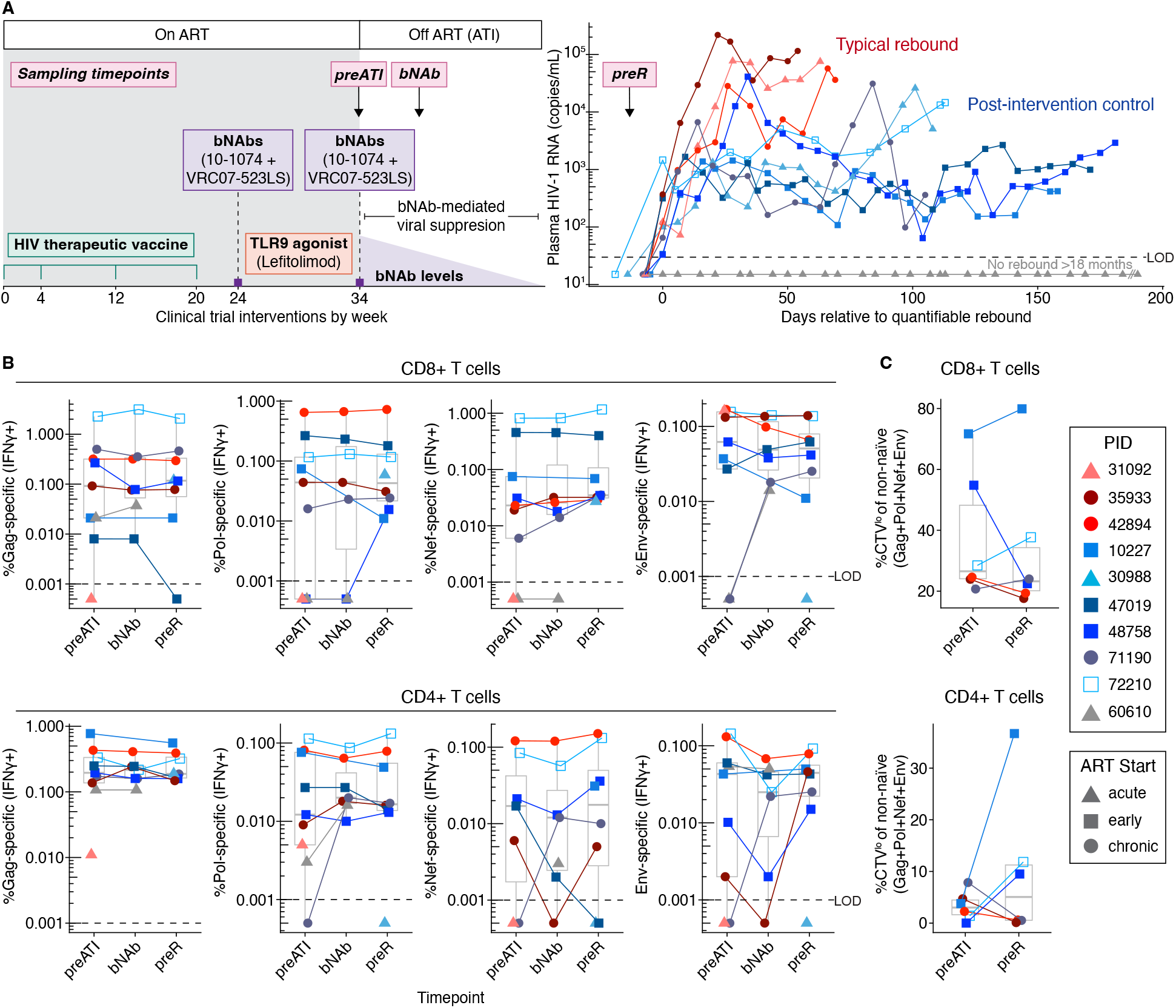
No evidence of enhanced HIV-specific T cell responses during bNAb-mediated HIV suppression. (A) Schematic of the clinical trial interventions and participant viral rebound curves with main sampling timepoints indicated (*preATI, bNAb, preR*); viral load curves end at the last timepoint prior to ART re-start. (B) Frequencies of Gag, Pol, Nef, and Env-specific CD4+ and CD8+ T cells at the *preATI, bNAb*, and *preR* timepoints as measured by intracellular cytokine staining for IFN*γ* in response to *in vitro* stimulation with relevant clade B overlapping peptide pools. (C) Sum of CD8+ or CD4+ T cell proliferative responses (Cell Trace Violet [CTV] dilution) in response to stimulation with HIV Gag, Pol, Nef, or Env clade B overlapping peptide pools at the *preR* vs *preATI* timepoints. Data were compared using linear mixed effect models with Benjamini-Hochberg false discovery rate correction (B) or Wilcoxon signed rank testing (C). No comparisons were significant (p or p_adj_<0.05). Box plots indicate median (line) and interquartile range (boxes).

### Stable HIV-specific T cell responses during bNAb-mediated HIV suppression

Because the vaccinal effect has classically been defined as a boost in antigen-specific T cell responses after mAb administration, we first investigated whether the magnitude and/or quality of HIV-specific T cell responses was enhanced during the period of bNAb-mediated viral suppression. Using standard flow cytometry-based *in vitro* peptide stimulation assays, we found no evidence at the *bNAb* or the *preR* timepoints that the magnitude or the proliferative responses of HIV-specific CD4+ or CD8+ T cells were boosted during bNAb-mediated suppression in our study, unlike the enhanced CD8+ T cell responses we had previously observed in response to rebounding virus (13) (**Fig. 1B, C**). More broadly, by cytometry time of flight (CyTOF), there was no significant increase in the frequency of total or non-naïve CD4+ or CD8+ T cells, or the frequency of activated (CD38+HLA-DR+) or cycling (Ki-67+) non-naïve CD8+ T cells (**Supp. Fig. 1A-C**). Finally, we looked at non-naïve CD8+ and CD4+ T cell clonotype expansion at the *preR* and *bNAb* timepoints compared to *preATI* by single cell (sc)RNA/T cell receptor (TCR)seq (268-1,927 [median 868] non-naïve T cells with TCR per sample) and only detected significant expansion of a single non-naïve CD8+ T cell clonotype in one participant at the *preR* timepoint (see **Supp. Fig. 1D** for representative analysis). Therefore, using a suite of T cell phenotyping, functional, and TCR repertoire profiling assays to evaluate samples early during bNAb-mediated HIV suppression and immediately prior to rebound, we did not find consistent evidence of the classical vaccinal effect, defined as activation or expansion of HIV-specific T cells, during bNAb-mediated HIV suppression in the combination immunotherapy trial.

### B cell activation during bNAb therapy in the absence of residual plasma viremia

Since the vaccinal effect has also been described as an enhancement of autologous antibody responses (5,6), we next looked at B cell activation and function during bNAb-mediated viral suppression after the ATI. Similar to T cells, there was no significant change in the frequencies of total or non-naïve B cells compared to *preATI* (**Fig. 2A**). However, we did observe a modest increase in the expression of the activation markers CD71 (transferrin receptor) and CD86 (T cell co-stimulation) on several B cell subsets at *bNAb* and/or *preR* compared to *preATI* (**Fig. 2B**,**C**). While we could not directly measure autologous anti-HIV antibody levels or functions due to interference from the bNAbs, we did measure polyclonal anti-HIV Env IgG3 levels (both bNAbs are the IgG1 isotype); there was no boost in these levels at *bNAb* or *preR* compared to *preATI* (**Fig. 2D**). Thus, while we observed no apparent increase in autologous antibody levels during bNAb-mediated suppression, there was modest evidence of B cell activation. Of note, this immune activation did not appear to be the result of stimulation of the cells by low-level viremia during bNAb-mediated HIV suppression, since essentially no residual plasma viremia was detectable using the Replicate Aptima single copy assay (SCA) during this period up until the *preR* timepoint (**Supp. Fig. 2, Supp. Table 3**) (25).

**Fig. 2:**
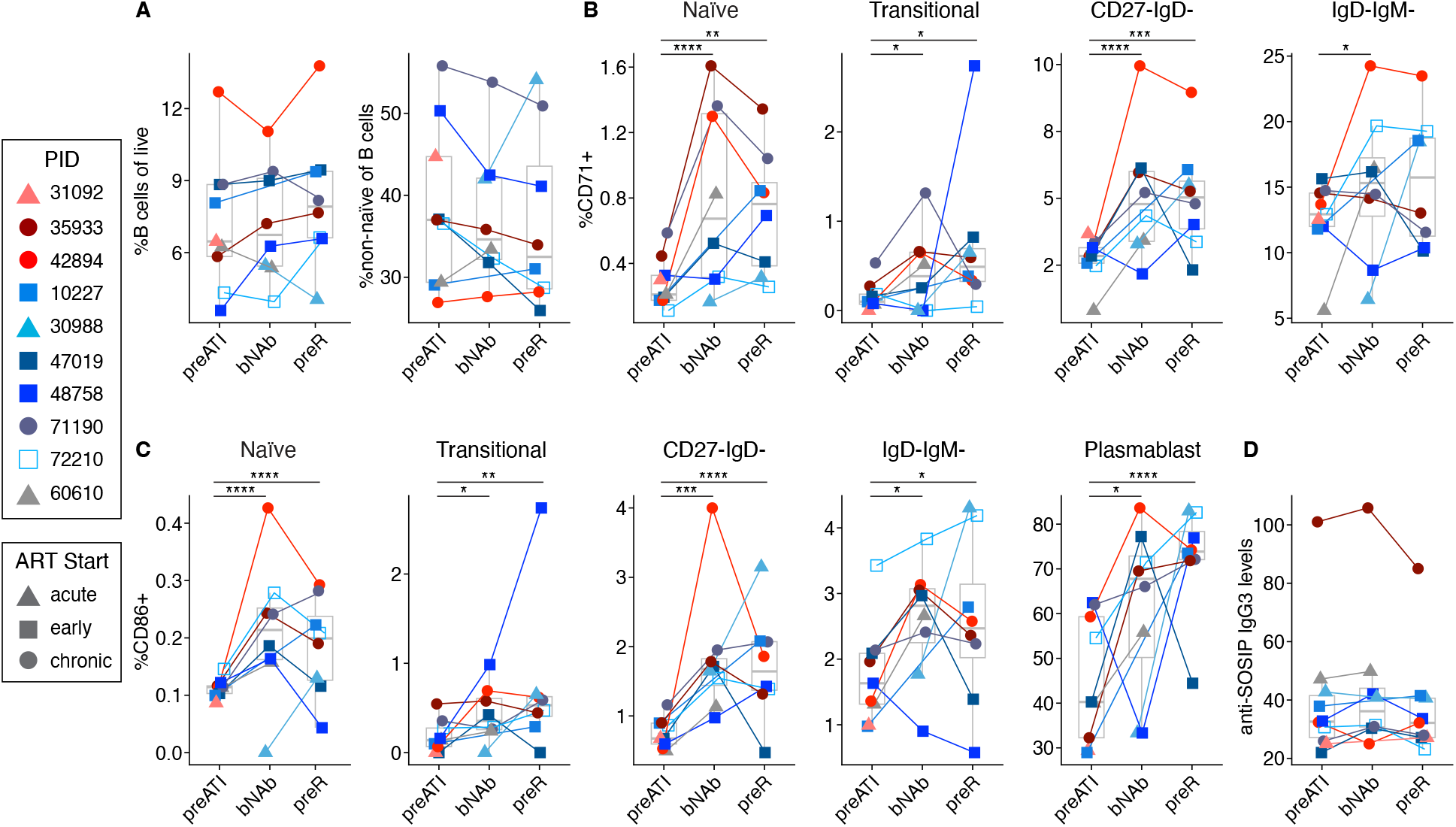
Modest changes in B cell activation during bNAb-mediated HIV suppression. (A) Frequencies of total and non-naïve B cells at the *preATI, bNAb*, and *preR* timepoints. Frequencies of CD71+ (B) or CD86+ (C) B cell subsets at the *preATI, bNAb*, and *preR* timepoints. (D) Plasma anti-SOSIP IgG3 levels at the *preATI, bNAb*, and *preR* timepoints. Data were compared using linear mixed effects models with Benjamini-Hochberg false discovery rate correction. * = p_adj_<0.05, ** = p_adj_< 0.01, *** = p_adj_<0.001, **** = p_adj_<0.0001. Box plots indicate median (line) and interquartile range (boxes).

### Broad innate immune cell activation during bNAb-mediated HIV suppression

We next evaluated changes in innate immune responses during the period of bNAb-mediated HIV suppression. Similar to adaptive immune cells, frequencies of major innate immune cell subsets were largely unchanged during the period of bNAb-mediated suppression, though there were modest, yet significant, increases in the frequencies of type 1 and 2 conventional dendritic cells (cDC1s, cDC2s) compared to *preATI* (**Fig. 3A**). Strikingly, however, several innate immune cell types demonstrated phenotypic evidence of activation as early as the *bNAb* timepoint, as measured using CyTOF. Specifically, at *bNAb* relative to *preATI*, we observed 1) a significant increase in the frequency of plasmacytoid dendritic cells (pDCs), cDC1s, and cDC2s expressing CD86 (**Fig. 3B**), 2) an increase in the proportion of pDCs and cDC2s expressing CD40, a protein involved in T and B cell co-stimulation (**Fig. 3C**), and 3) an increase in the proportion of cDC1s and cDC2s as well as classical (CD14+ CD16-) and intermediate (CD14+CD16+) monocytes expressing CD71 (**Fig. 3D,E**). These activation markers generally remained elevated at the *preR* timepoint relative to *preATI* as well. In contrast to other innate immune cell types, NK cells did not show consistent evidence of activation during bNAb-mediated suppression, though there was a significant increase in the proportion of Perforin+ CD56^hi^ (more immature) NK cells at *preR* relative to *preATI* (**Supp. Fig. 3**). Taken together, these data demonstrate that several innate immune cell types important for viral sensing and cytokine production as well as antigen presentation and co-stimulation of adaptive immune cells become activated early after bNAb administration, weeks before quantifiable rebound occurred. Of note, based on the phenotypic markers we characterized, innate immune activation was similar between post-intervention controllers and non-controllers.

**Fig. 3.**
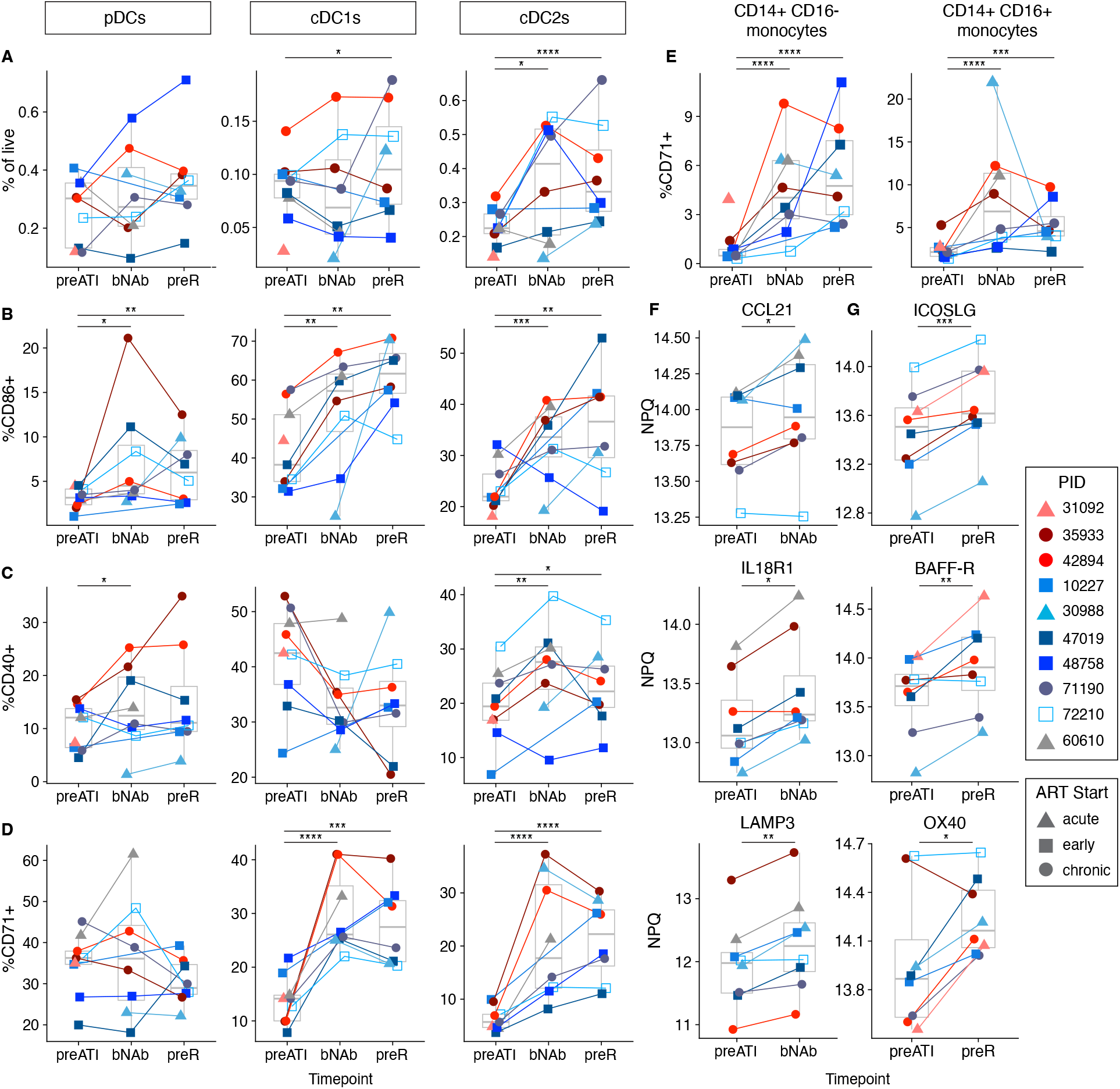
Evidence of innate immune activation during bNAb-mediated HIV suppression. (A) Frequencies of indicated innate immune cell subsets as a proportion of live CD45+ cells by CyTOF at the *preATI, bNAb*, and *preR* timepoints. Frequency of CD86+ (B), CD40+ (C), or CD71+ (D, E) of the innate immune cell subsets indicated at the indicated timepoints. Log_2_ normalized plasma protein (NPQ) levels as measured by NULISAseq at the *preATI* versus *bNAb* (F) or *preATI* versus *preR* (G) timepoints. Data were compared using linear mixed effects models with Benjamini-Hochberg correction (A-E) or paired t-tests without false discovery rate correction (F, G). * = p or p_adj_<0.05, ** = p or p_adj_< 0.01, *** = p or p_adj_<0.001, **** = p or p_adj_<0.0001. Box plots indicate median (line) and interquartile range (boxes).

### Plasma inflammatory proteins increase during bNAb therapy

We next asked whether the changes we observed in innate immune cell activation during bNAb-mediated HIV suppression were accompanied by changes in plasma inflammatory protein levels. To do this, we compared protein levels at the *preATI* versus *bNAb* and *preR* timepoints using Nucleic acid Linked Immuno-Sandwich Assay (NULISAseq), a highly sensitive multiplexed immunoassay that detects a panel of 250 immune-relevant plasma proteins (26) (see **Supp. Table 4** for full dataset). No plasma protein levels were significantly changed at *bNAb* versus *preATI* or *preR* versus *preATI* using t-tests with a stringent false discovery rate correction (p_adj_<0.05). However, we noted that a subset of plasma proteins that passed an unadjusted p-value significance threshold (p<0.05) subtly increased in nearly all participants after the ATI and before rebound. Already at the *bNAb* timepoint a month prior to quantifiable rebound, inflammatory proteins like CCL21, IL18R1, and LAMP3 were increased relative to *preATI* (**Fig. 3F**). These proteins help bridge the innate and adaptive immune responses, promoting dendritic cell antigen presentation as well as lymphocyte trafficking and differentiation. Additional inflammatory proteins directly involved in promoting T and B cell activation and differentiation, including ICOSLG (ICOS ligand), were increased at the *preR* timepoint compared to *preATI* (**Fig. 3G**). These data suggest that increases in immune-relevant inflammatory plasma proteins may occur under bNAb-mediated HIV suppression, well before quantifiable viremia emerges.

### Transcriptional activation of immune cells during bNAb-mediated HIV suppression

We next performed CITEseq (27), which combines scRNA/TCRseq with surface protein expression, on PBMCs isolated from the *preATI, bNAb, preR*, and *postR* timepoints to more comprehensively characterize the transcriptional pathways that become activated in immune cells during bNAb-mediated viral suppression (see **Fig. 4A** for broad cell type annotation; see **Supp. Figs. 4** and **5** for higher resolution cell type subclusters and genes and proteins used to identify them). Mirroring our CyTOF data, the frequency of most broad immune cell clusters did not change consistently across participants during the period of bNAb-mediated viral suppression after the ATI and prior to rebound (see **Supp. Fig. 6** and **7** for cluster frequencies over time). Using a stringent statistical cut-off, there were very few (1, 2, or 32 at p_adj_<0.1) differentially expressed genes within the cell type subclusters across timepoints. In contrast, using gene set enrichment analysis (GSEA) to identify groups of genes that changes over time within each cell type subcluster between *preATI* and *bNAb*, we found that the expression of genes within early viral response pathways such as IFNα, TNFα, and IL-6 signaling as well as antigen cross-presentation was significantly enriched as early as the *bNAb* timepoint in several innate immune subclusters, particularly in cDC2s (**Fig. 4B**). Many of the genes in these pathways are important for antiviral sensing and coordination of innate and adaptive immune responses (e.g., *RIGI, IFITM3, TAP1, CCL20, CXCL10, IFNGR2, IL18, IL1A, IL1B, CD80*; **Fig. 4C, D**). These pathways generally became further enriched at the *preR* timepoint. While changes in early viral response pathways post-ATI were mostly restricted to innate immune cell types, we also observed an enrichment in the expression of genes in translation and Myc signaling pathways mostly in adaptive immune cell subclusters at the *bNAb* and/or *preR* timepoints relative to *preATI* (**Fig. 4B**). For both the innate and adaptive immune cell subclusters, while the expression of genes within these select pathways was more pronounced in some participants compared to others, upregulation generally occurred consistently between post-intervention controllers and non-controllers (**Supp. Fig. 8**).

**Fig. 4.**
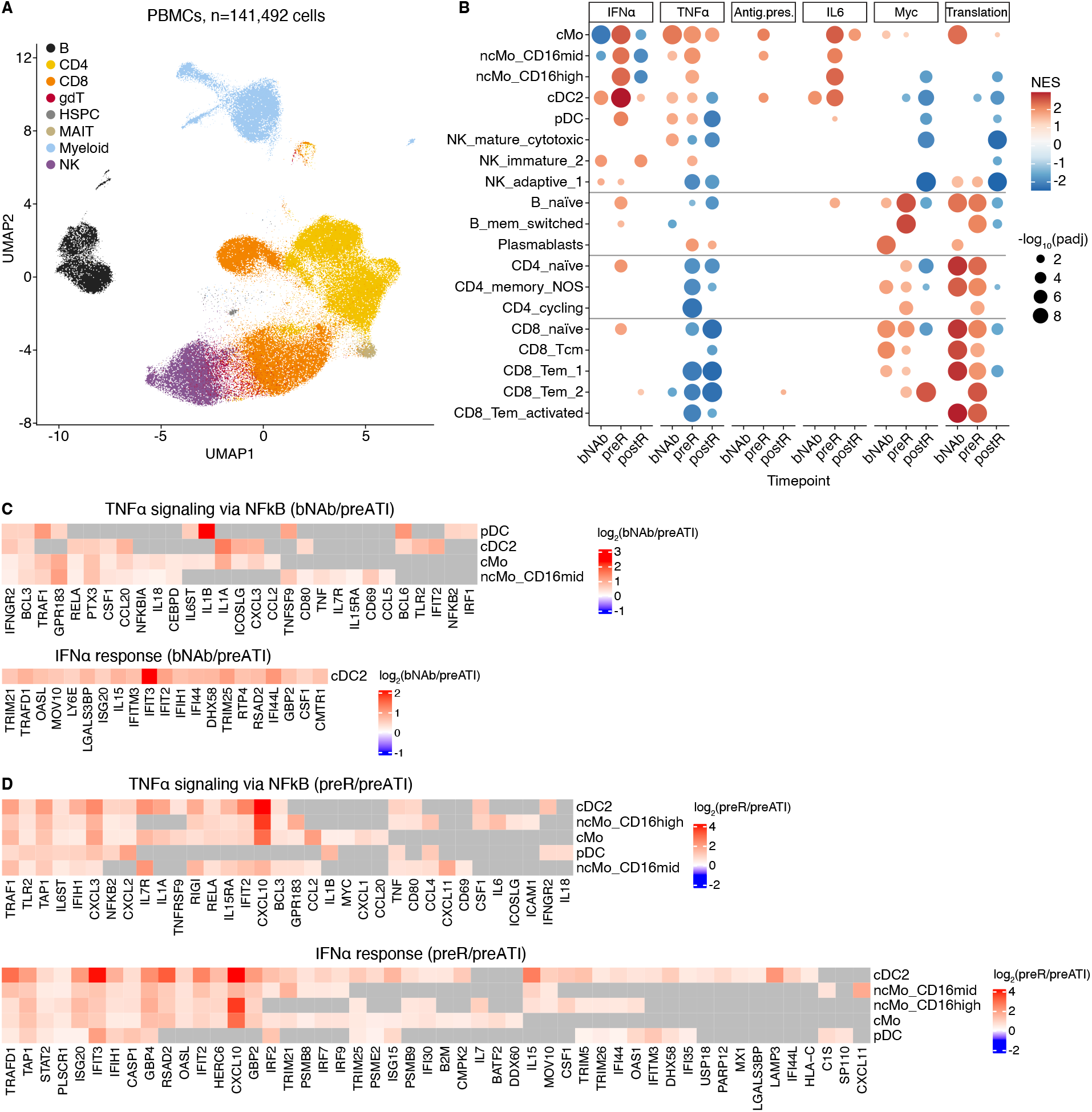
Transcriptional activation of innate and adaptive immune cells during bNAb-mediated HIV suppression. (A) UMAP of broad PBMC annotations by CITEseq (see **Supp. Figs. 4-5** for cell type subclusters). (B) Gene set enrichment analysis bubble plots depict normalized enrichment scores (NES; relative to *preATI*) for the pathways listed in the indicated cell subclusters at the indicated study timepoints. Pathways include Hallmark Interferon (IFN)α response, Hallmark TNFα signaling via NFκB, Reactome Antigen presentation folding assembly and peptide loading of class I MHC, Hallmark IL6 JAK STAT3 signaling, Hallmark Myc targets V1, and Reactome translation. Red indicates positive enrichment and blue negative enrichment. Bubble size corresponds to the negative log_10_ of the adjusted p-value. Select leading-edge genes in the IFNα response and TNFα signaling pathways at the (C) *bNAb* versus *preATI* and (D) *preR* versus *preATI* timepoints. Heatmaps denote log_2_ fold change in gene expression relative to *preATI*. Grey indicates no data.

### Unique patterns of immune activation in post-intervention controllers

Although several markers of immune activation during bNAb-mediated HIV suppression appeared similar between post-intervention controllers and non-controllers in the combination immunotherapy trial, we observed several important differences between the two groups. Applying GSEA to our CITEseq differential expression data, we observed that at the *preATI* timepoint, genes in the Hallmark TNFα and inflammatory response signaling pathways were enriched in innate immune cell types (conventional monocytes, cDC2s, pDCs) of post-intervention controllers compared to non-controllers (**Fig. 5A, Supp. Fig. 9**). Genes in these pathways were similarly enriched in innate immune cells from post-intervention controllers at the *bNAb* timepoint (**Fig. 5A**). Many of the enriched genes play key roles in innate immune cell activation and effective priming of the adaptive immune response (e.g., *ICOSLG, CD80, TNFAIP3, IL1B, CCL4, TNF, IL6, CXCL10*, and *CCL20*; **Fig. 5B**). While the TNFα pathway was not differentially active between controllers and non-controllers in adaptive immune cell subclusters at the *preATI* or *bNAb* timepoints, controllers demonstrated greater enrichment across several adaptive immune cell types (e.g., memory B cells that have not undergone isotype switching, naïve CD4+ and CD8+ T cells) at the *preR* timepoint (**Fig. 5A, Supp. Fig. 9**). Here, we observed enrichment of genes involved in lymphocyte activation (e.g., *JUN, FOS*, and *NFAT5*; **Fig. 5B**). Additional differences between post-intervention controllers and non-controllers emerged when we modeled the NULISAseq plasma proteomics data at multiple timepoints post-ATI. First, while expression of plasma proteins involved in adaptive immune activation/co-stimulation (e.g., CD80 and IL12B) were relatively stable following ATI in post-intervention controllers, these tended to be lower during the period of bNAb-mediated viral suppression in non-controllers, with levels catching up with or surpassing those of controllers after viral rebound (**Fig. 5C**). Second, type I interferons (e.g., IFNA2, IFNA1/13, and IFNW1) were relatively stable following ATI in post-intervention controllers; however, levels of these proteins increased dramatically after viral rebound in non-controllers, and indeed appeared to begin increasing even prior to overt rebound (i.e., at the *preR* timepoint, **Fig. 5D**). Together, these data may point to an immune environment that is less conducive to adaptive immune priming during bNAb-mediated viral suppression in non-controllers, which may result in an impaired ability to control even the earliest reactivating virus in tissues.

**Fig. 5.**
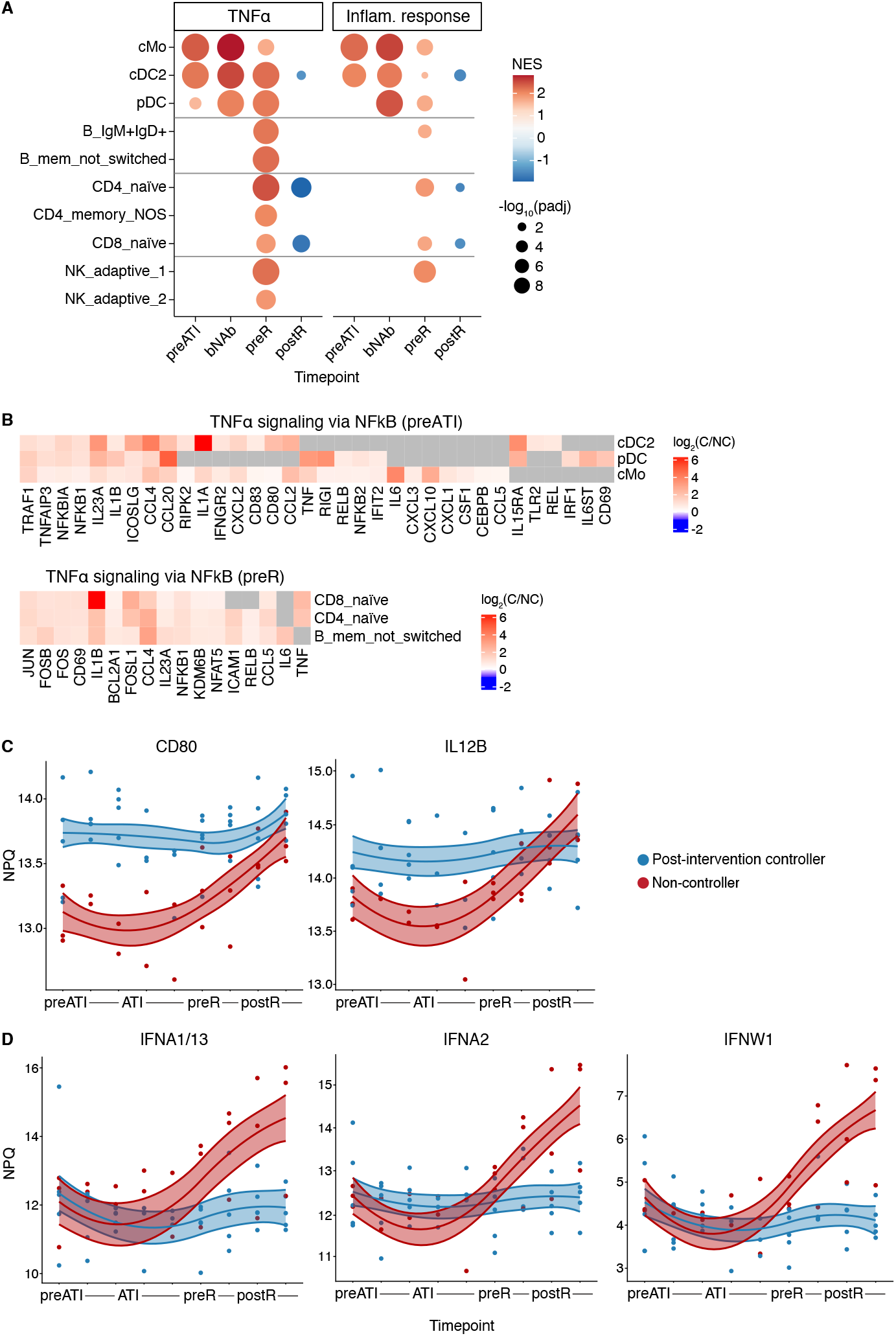
Differential immune activation between post-intervention controllers and non-controllers. (A) Gene set enrichment analysis bubble plots depict normalized enrichment scores (NES; post-intervention controllers relative to non-controllers) for Hallmark TNFα signaling via NFκB and Inflammatory response pathways in the indicated cell type subclusters at the indicated study timepoints. Red indicates positive enrichment and blue negative enrichment. Bubble size corresponds to the negative log_10_ of the adjusted p-value. (B) Select leading-edge genes in the TNFα signaling pathway at the indicated timepoints in the indicated cell types. Heatmaps denote log_2_ fold change in gene expression in controllers relative to non-controllers. (C, D) Spline models of plasma protein levels over time showing significant differences between controller status (in blue post-intervention controller and in red non-controller participants) at p_adj_<0.05.

## DISCUSSION

Several studies have shown that bNAbs administered during periods of HIV antigen expression can promote some degree of viral control in up to 30-40% of participants after bNAb levels wane, however the mechanisms underlying this effect remain unclear and no specific bNAb-induced immune responses have yet been linked to post-bNAb control (9–13,19,28–34). In this analysis of a combination immunotherapy trial, we found evidence of immune activation and enhancement during bNAb-mediated HIV suppression following ATI using multiple complementary approaches. Specifically, we found clear evidence of innate and adaptive immune cell activation that occurred early after ATI/bNAb infusion, more than a month prior to quantifiable rebound. Furthermore, while much of this activation occurred similarly across study participants, we also identified differences in several patterns of immune activation between post-intervention controllers and non-controllers during this critical window.

This report represents one of the first descriptions of innate immune activation occurring under bNAb-mediated HIV suppression, within a few weeks of stopping ART and as early as one month prior to quantifiable rebound. We observed discrete activation of specific innate immune cell types, most notably dendritic cells (pDCs, cDC1s, cDC2s) and monocytes (both classical and intermediate), as evidenced by upregulation of activation/co-stimulation markers, genes in early viral responses pathways, and inflammatory plasma proteins under bNAb-compared to ART-mediated HIV suppression. Some of the innate immune activation that we observed would be expected to result in the stimulation of adaptive immune responses (e.g., the upregulation of CD40 and CD86 on antigen presenting cells). Notably, we also observed upregulation of genes in Myc and translation signaling pathways in a subset of adaptive immune cells as well as modest B cell activation (as evidenced by upregulation of CD71 and CD86) during the period of bNAb-mediated HIV suppression. This immune activation occurred in the absence of residual low-level viremia.

Enhanced immune activation during bNAb-mediated HIV suppression may have important implications. Mechanistically, we hypothesize that the immune activation we observed after bNAb infusion played a role in enhancing subsequent HIV-specific adaptive immune responses. Indeed, while much of the broad immune activation we observed occurred similarly across participants, there were some important distinctions between post-intervention controllers and non-controllers. Even prior to the ATI, innate immune cells of post-intervention controllers had significantly higher expression of genes in the TNFα and inflammatory response pathways, including transcripts for several genes important for sensing and responding to viral infections; these gene expression differences persisted into the period of bNAb-mediated HIV suppression. Additionally, immediately prior to rebound, presumably just as immune responses were preparing to respond to the burst of rebounding virus, post-intervention controllers had higher expression of TNFα response pathway genes important for lymphocyte activation in several adaptive immune cell types compared to non-controllers. Controllers also had consistently higher levels of plasma proteins involved in the stimulation of adaptive immune cells (e.g., IL12B and CD80) during the period of bNAb-mediated viral suppression. Just prior to and into the earliest phases of rebound, while controllers and non-controllers similarly engaged several pathways of immune activation, non-controllers demonstrated a more profound increase in type I interferon levels in the plasma. Taken together, these data indicate that, during bNAb-mediated HIV suppression and long before viral rebound, post-intervention controllers had innate immune cells that were more poised to respond to reactivating virus and effectively present antigen and provide co-stimulation to adaptive immune cells, which in turn were able to more effectively become transcriptionally activated as the virus was emerging, even prior to overt rebound.

We did not observe direct enhancement of HIV-specific adaptive immune responses during bNAb-mediated HIV suppression, which may have been explained by several factors: (1) we evaluated post-bNAb responses too early: other bNAb studies describing a vaccinal effect sampled roughly 8-12 weeks post-ATI (9,10,12,14,21) while our bNAb timepoints was ~4 weeks into the ATI (although we did not observe augmentation of HIV-specific adaptive immune responses at the *preR* timepoint either), (2) enhanced responses may only be found in tissues at sites of viral persistence and not in the blood, and/or (3) we needed to measure different functions or specificities (e.g., activity against autologous virus) in order to capture the relevant T cell and/or antibody responses. Ongoing and future planned studies will address these limitations directly.

Other studies have described similar immune activation following cessation of ART and prior to rebound. In one, upregulation of antiviral and inflammatory signaling pathways was observed by bulk RNAseq during bNAb-mediated suppression and prior to viral rebound; these changes were not associated with post-intervention control and the relative contribution of individual immune cell types to the shift in transcriptional signature was not delineated (35). Other studies have described a similar phenomenon in the absence of bNAb infusion or any immune-based interventions. Specifically, activation of pDCs after interruption of ART has been shown to correlate with time to viral rebound (36), and upregulation of antiviral transcriptional pathways and expansion of CD16++ non-classical monocytes has been described in prior viremic controllers following ATI (35,37). These data suggest that an enhanced ability to detect rebounding virus and activate innate immune responses may be important for post-ART viral control even in the absence of an immunotherapy. Our study adds to this literature by deeply characterizing immune activation using sensitive single cell and plasma assays under bNAb-mediated suppression.

Our study had important limitations. It was a small, single-arm, proof-of concept study with 10 participants. The study also did not include a placebo arm, which limited our ability to clearly distinguish bNAb-mediated changes from those that may have been driven by the other immune interventions that participants received (e.g., the therapeutic vaccine). In light of the sampling timepoints investigated, the immune activation we observed under bNAb-mediated suppression may also have been at least in part due to non-specific bNAb-(as opposed to bNAb:HIV immune complex)-mediated immune activation. The fact that we observed differential immune activation between controllers and non-controllers, however, argues against this immune activation being entirely non-specific. Finally, given the increasing use of bNAbs in therapeutic settings (38), the immune activation we occurred during bNAb-mediated suppression could have clinical implications, but would need to be studied in larger cohorts and over time to understand long-term consequences.

In sum, we provide evidence that broad immune activation occurs during bNAb-mediated HIV suppression and identify candidate immune pathways that become activated in this window and may be linked to post-intervention control of HIV. Future evaluation of these pathways in larger clinical studies of people with HIV who receive bNAbs, including a close evaluation of tissue samples to monitor immune responses, the reservoir, and bNAb pharmacokinetics, will be required to parse these mechanisms.

## METHODS

### Study cohort

For full details of the study cohort, please see (13). Briefly, the UCSF-amfAR combination immunotherapy trial (NCT 04357821) enrolled 10 people with HIV. 3 participants had initiated ART during acute HIV infection (<30 days after acquisition), 4 during early HIV infection (between 1 and 6 months after HIV acquisition), and 3 during chronic HIV infection (≥6 months after HIV acquisition). Participants were required to have been on suppressive ART for over 12 months, to have a screening CD4+ T cell count ≥500 cells per uL, and to lack exclusionary comorbidities. Participants whose reservoir provirus was determined to have significantly reduced phenotypic susceptibility to one or both bNAbs were excluded. The study was approved by the UCSF IRB. Sampling timepoints for this study (*preATI, bNAb, preR*) were selected as above. Two participants (PID 10227 and 31092) did not have PBMCs available at the *bNAb* timepoint, and two (60610 and 31092) at the *preR* timepoint.

### Peripheral blood mononuclear cell and plasma isolation and storage

At most study visits, peripheral blood was collected by standard blood draw or leukapheresis and stored as serum, plasma, and viably cryopreserved PBMCs, as described previously (39). Plasma was stored at −80°C and PBMCs were stored in liquid nitrogen.

### Clinical viral load measurements

As previously described (13), plasma HIV-1 RNA levels were initially measured using the Abbott Real Time HIV-1 PCR assay (limit of quantitation 40 copies/mL) and subsequently with the Hologic Aptima assay (limit of quantitation, 30 copies/mL). Plasma HIV RNA levels <30 copies/mL were considered unquantifiable. Laboratory measurements were performed in the clinical laboratory at Zuckerberg San Francisco General Hospital.

### Replicate Aptima assay for ultrasensitive quantification of residual plasma viremia

Residual plasma HIV-1 RNA was quantified by the Vitalant Research Institute (VRI) using a replicate testing strategy with the Aptima® HIV-1 Quant Dx assay (Hologic, Inc.) on the fully automated Panther® system, as developed and validated at VRI (40,41). By running multiple replicates on a fully automated commercial platform, this method offers a scalable, high-throughput alternative to manual single-copy assays. Up to nine replicate tests, each processing 500 µL of plasma, were performed for each sample. For each replicate, the Panther® System reports one of three outcomes: a quantitative result (≥30 copies/mL), a detected-but-not-quantified signal (<30 copies/mL), or no signal detected. Replicates were classified as positive (either of the first two outcomes) or negative (third outcome), and the resulting positivity rate was used to estimate viral concentration via a Poisson statistical model, with a correction factor applied to account for imperfect per-replicate sensitivity (40,41). When at least one negative replicate was present, 95% confidence intervals (CIs) were calculated from the Poisson-based maximum likelihood estimate as previously described (42). In cases where all replicates were positive, which precludes Poisson-based estimation, viral concentration was estimated as the arithmetic mean of per-replicate values, where replicates with a detected-but-not-quantified signal (<30 copies/mL) were assigned a concentration using Hologic’s proprietary extrapolation algorithm. In this situation, 95% CIs were calculated as 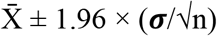, where 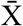 is the mean, ***σ*** the standard deviation, and n the number of replicates.

### Modeled broadly neutralizing antibody levels

For methods related to the quantification and modeling of bNAb levels, see (13).

### Antibody isotype and subclass binding

HIV-antigen specific antibody isotype and subclass titers were measured in plasma samples via Luminex multiplexing, as previously described (43). In short, JRFL SOSIP (Duke University Protein Core, USA) antigens were directly coupled to distinct Luminex bead regions (Luminex Corp, TX, USA) through Sulfo-NHS (Thermofisher) and EDC (Thermofisher) coupling. Plasma samples were diluted (1:50 for isotype and subclass) and added to the coupled bead mix in 384 well plates (Greiner Bio-One, Germany) for a 2-h incubation at room temperature. excess antibodies were washed away. Antibody isotypes were stained with mouse anti-human IgG-PE antibodies (Southern BioTech) for 1 h at room temperature. Plates were washed again, and beads were resuspended and run on flow cytometry instruments (Luminex INTELLIFLEX) to determine the geometric mean fluorescent intensity. All experiments were run in duplicate fashion.

### NULISAseq

Plasma samples were stored at −80°C until ready to use, then thawed and centrifuged at 10,000g for 10min. 10μL of supernatant for each sample was plated in a randomized order in a 96 well plate and assayed using the NULISAseq Inflammation Panel 250 (26). Samples were run on two plates with assay controls on each plate (sample controls, inter-plate controls, and negative controls) and an internal control (exogenous reporter) was added to each well. The automated Alamar ARGO™ prototype system was used for immunocomplex formation with the paired set of oligo-conjugated antibodies, first capture with oligo-dT beads, release, and second capture with streptavidin beads and ligation. The library of DNA reporters containing unique target-specific molecular identifiers and sample-specific molecular identifiers was pooled, amplified by PCR, purified, and sequenced on the Illumina NextSeq 2000 system. Data were normalized using internal control-based normalization then divided by target-specific inter-plate control medians; data were then rescaled by multiplication with a factor of 10^4^. Finally, +1 was added to each value and data were then log_2_ transformed, generating NULISA Protein Quantification (NPQ) units used for data analysis. For quality control, internal control counts were required to be within +/-40% of the median internal control counts across the plate. 12/170 samples did not pass quality control, all of which came from a single participant (48758). Due to the possibility of matrix effects and interference due to specific biological factors, this participant’s samples were excluded from downstream analyses.

### NULISAseq between-timepoint statistical comparisons

Comparisons of protein levels between timepoints (*bNAb* or *preR* comparted to *preATI*) were performed in R (v4.5.0) (44) using paired t-tests. There was not a significant difference in protein levels when Benjamini-Hochberg false discovery rate correction was applied, we explored proteins whose levels differed between timepoints at an unadjusted p<0.05.

### NULISA spline models

To determine differential changes in plasma protein levels across time between controllers and non-controllers, we used spline models for their flexibility, allowing us to capture different profiles of changes before and after viral rebound. Results were similar with more standard quadratic models. We included data from the following timepoints (median days relative to ATI [interquartile range]): *preATI* (−2 [−2 to −2]), the first PBMC timepoint collected after the ATI (*ATI1*; 11 [11 to 26.5]), *ATI2* (25 [24.5 to 39]), *ATI3* (43 [40 to 66]), *ATI4* (67 [55 to 82]); and the following timepoints (median days relative to rebound [interquartile range]): *preR* (−10.5 [−19.25 to −6.75]), *postR1* (8 [6.75 to 13.25]), *postR2* (25 [21.75 to 35]), and *postR3* (42 [37.5 to 53]). We used B-spline mixed effects models to detect smooth, non-linear NULISA protein target trajectories over time (*preATI* to *postR3*). We encoded the timepoint as an ordinal variable ranging from 0 (*preATI*) to 8 (*postR3*). We used the bs()function implemented in splines R package (45) to estimate B-spline functions over this interval setting and internal knot at the pre-rebound timepoint and a quadratic trend (degree = 2). The basis functions were generated using the Cox–de Boor recursion algorithm. Given these parameters, the resulting B-spline representation consists of three basis functions, each contributing differentially across the time, with the first basis function dominating at early time points and the third basis function contributing most strongly at later time points. Smooth temporal profiles were modeled as linear combinations of these three basis functions. Specifically, different curve shapes were obtained by varying the coefficients assigned to each basis function, enabling flexible representation of diverse temporal patterns. For each target, model fitting using lmer() involved estimating the optimal set of coefficients for the three basis functions that best captured the smooth nonlinear relationship of the observed NPQ values over time, where participant ID was modeled as random effect to account for within-individual correlation. This approach is analogous to fitting a quadratic model, where the response is expressed as a linear combination of the basis functions (1), (t), and (t^2), and the coefficients are estimated to best fit the observed data. We compared this model to a model that included the spline by-status interaction term using an ANOVA test to evaluate whether including the status improved the fit. The p-values were adjusted for multiple comparisons using Benjamini-Hochberg false discovery rate correction. For visualization, we estimated the population mean NPQ values separately for controllers and non-controllers at 0.2 time-intervals between *preATI* and *postR3* using the fixed effects and variance-covariance matrix from the model fits. We visualized the observations, predicted population mean, and standard error curves using the *gpplot2* R package (46). In addition to the participant 48758 excluded for low quality of the NULISAseq data, participant 60610 was excluded from spline models because post-rebound timepoints were not collected.

### CITEseq library preparation

PBMCs were thawed, counted, and combined (15k cells per sample, 3-4 samples/batch) and loaded onto 10x chips. Single-cell expression libraries were prepared using the 10x Genomics Chromium Next GEM Single Cell 5’ GEM Kit v2 and Library Construction Kit, according to the manufacturer’s protocol. Briefly, 45,000-60,000 cells were loaded into the 10x Chromium Next GEM Chip K and run on the 10x Chromium Controller to generate Gel Beads-in-emulsion (GEM), followed by incubation to produce barcoded, full-length cDNA from polyadenylated mRNA (5’ UTR to constant region) while simultaneously capturing the cell surface protein sequence. The amplified cDNA from polyadenylated mRNA was used to generate gene expression, T-cell receptor, and B-cell receptor libraries, and the amplified DNA from cell surface protein Feature Barcode was used to generate cell surface protein libraries. Following quality control of libraries, a pool of the libraries was sequenced on the Illumina NovaSeq X 25B flow cell.

### CITEseq data pre-processing, demultiplexing, doublet detection, and quality control

Sequencer-obtained raw FASTQ data were aligned to the GRCh38 reference genome (GENCODE v44/Ensembl110) to obtain feature-barcode matrices for each library and the raw FASTQ data of surface protein expression to the TotalSeq™-C antibody reference panel (**Supp. Table 5**) using the Cell Ranger count (v8.0.0; multi command). Raw feature-barcode matrices were imported and analyzed using Seurat (v5.1.0) (47).

Each library was demultiplexed to map individual cells to their sample of origin using freemuxlet (vAug2021) (48), SNP profiles were mapped to participants’ genotypes from bulk RNA-seq using bcftools gtcheck (v1.10.2) (49), and inter-sample doublets and empty droplets were removed. DoubletFinder (v2.0.3) (50) was used to filter barcodes containing heterotypic intra-sample doublets, as described (51), with default parameters except as specified: npcs=35, expected rate of intra-sample doublets = inter-sample doublet rate as estimated by freemuxlet divided by (*number of pooled samples – 1*). Barcodes containing fewer than 100 genes and genes that are detected in fewer than 3 cells were removed. We further filtered out the barcodes based on library-specific cutoffs for the number of genes per cell (nFeature_RNA <600-1100 and >5,000-6,000), the number of unique molecular indexes (UMIs) per cell (nCount_RNA <1,000-3,000 and >20,000-25,000; nCount_ADT >20,000), mitochondrial content (>12%), ribosomal content (<8% and >60%) to remove poor quality cells. We additionally removed cells in which the top-expressed gene accounted for more than 70% of detected molecules. Furthermore, cells identified as platelets (defined as >1 UMI for PPBP, PF4, CAVIN2, and/or GNG11 genes) or red blood cells (>1 UMI for HBB, HBA1, and/or HBA2 genes) were excluded. The resulting dataset of 170,255 high-quality cells was used for further analysis.

### CITEseq data normalization, dimensionality reduction, and clustering of major cell types

Gene expression data from all libraries were merged, log-normalized, and scaled with regression on UMI and gene counts, mitochondrial content, ribosomal contents, and predicted cell-cycle state. The top 2,000 variable features were identified using the “vst” method, TCR genes were excluded from the variable features, PCA was performed using npcs=50, and batch-effect (library-specific bias) was normalized using Harmony (52). ADT data were normalized and denoised using the “denoised and scaled by background” (dsb) (53) method for each library separately using empty droplets as background. The empty droplets were defined, separately for each library, as barcodes with fewer than 100-350 UMIs and 10-55 < ADT counts < 100-550. The signal from isotype-control antibodies was used to denoise the data. Normalized values less than −10 were truncated. All ADTs, except for those corresponding to isotype control antibodies, were used as variable features. We used reciprocal PCA to batch-correct and integrate the protein data. Briefly, after scaling the data and calculating principal components (PC) for each library, the data was integrated using the IntegrateData function in Seurat. The integrated protein data was scaled, the ADT counts per cell and cell-cycle states were regressed out, and PCA was performed using npcs=30.

A weighted nearest neighbor (WNN) graph was constructed using the first 30 harmony dimensions for gene expression and the first 18 integrated PCs for protein expression using Seurat’s FindMultiModalNeighbors function with default parameters except prune.SNN = 1/20. Using the WNN graph, dimensionality reduction was performed with Uniform Manifold Approximation and Projection (UMAP), and single-cell clustering with the smart local moving (SLM) algorithm in Seurat’s FindClusters function. We generated clusters using resolution values between 0.5 and 4 and, based on manual inspection, selected 0.6 to obtain 26 cell clusters.

Upon manual inspection of individual clusters, we identified two clusters (9 and 12) that contained cells from two or more different cell types, namely small subsets of CD4+ T cells, CD8+ T cells, and NK cells, which we separated by an additional round of clustering of the two clusters separately, resulting in a total of 30 cell clusters. We grouped these clusters into 7 major cell types, including B, CD4+ T, CD8+ T, other T (clusters of MAIT, *γ*δT, Treg, and proliferating T cells), myeloid, NK, and Hematopoietic Stem and Progenitor Cells (HSPCs) based on gene and ADT expression profiles. Three clusters that lacked unique marker expression, comprising 12,664 cells (7.4%), were discarded. Consistent with previously reported (54) difficulty with unsupervised clustering in clearly separating CD4+ T and CD8+ T subsets, we observed CD4 ADT expression in CD8+ T cell clusters and vice versa. Moreover, a small number of T cells also expressed canonical surface markers of B and myeloid cells, suggesting the presence of residual doublets. To identify pure CD4+ and CD8+ T populations, we combined CD4+, CD8+, and other T populations and manually selected T cell subsets based on expression of their canonical markers in RNA and ADT data. T cells were defined as CD3+ (ADT > 25 or RNA > 0.5 for at least one isoform), CD19-(ADT < 45), CD20-(ADT < 38), CD14-(ADT < 32), and CD11c-(ADT < 45). Among them, the cells expressing CD4 (ADT > 30 or RNA > 0.5) were classified as CD4+ T, and cells expressing CD8 (ADT > 32 or RNA > 0.5 for at least one isoform) as CD8+ T cells. Cells that did not satisfy these selection criteria (9,727; 5.7%) were excluded from the analysis. Cutoffs were defined based on the distribution of marker expression and the corresponding marker-positive populations.

### Identification of cell subsets via subclustering

Each major population, including B cells, NK cells, myeloid cells, CD4+ T cells, and CD8+ T cells, was subclustered to identify cell subsets within each compartment. The same procedure for data normalization, batch correction, and clustering as described above was used for the subclustering, with the following modifications for T cells. Through a manual inspection of ADT expression in the T cell compartment, we identified markers with high background/non-specific expression or those that are irrelevant for T cells (n=41) (**Supp. Table. 5**). To avoid introducing noise from these markers, we excluded them from the set of highly variable features, along with the ADT markers used to manually select CD4+ T cells and CD8+ T cells. Our preliminary analysis of T cell subsets revealed a recognizable effect of proliferation on clustering. To avoid the effects of proliferation on cell type identification, genes correlating (Spearman’s correlation >0.2) with either *MKI67, TOP2A, H2AC11*, or *H1-4* were excluded from the set of highly variable genes at the start, as described previously (55).

We generated clustering using resolution values between 0.3 and 1 and evaluated them using clustree (56). The resolution that provided the greatest separation of functionally distinct subpopulations was selected for each major population. The markers that distinguish individual clusters were identified using Model-based Analysis of Single Cell transcriptomics (MAST) (57) in FindAllMarkers function of Seurat (test.use = “MAST”, max.cells.per.ident =1000, random.seed = 1234, and latent.vars = “batch”) and were used to annotate clusters (**Supp. Fig. 5**). Normalized gene expression and centered log-ratio (CLR)-transformed data were used to identify gene and protein markers, respectively. After excluding small clusters without distinct marker expression (6,372; 3.7%), we identified 42 distinct immune cell subsets, comprising 141,492 cells.

Samples corresponding to the second post-rebound timepoint for all participants were excluded from the downstream analyses because they were outside the scope of the analysis, which focused on the period of bNAb-mediated suppression and early into viral rebound. The first post-rebound timepoint for participant 35933 was excluded from the downstream analyses due to significant plasma viremia at this timepoint (>2600 copies/mL). No samples beyond bNAb for participant 60610 were included in the downstream analyses since no quantifiable rebound was detected for this participant throughout the duration of study monitoring.

### Differential gene expression analysis and pathway enrichment analysis

To identify gene expression changes over time after ATI, we compared each timepoint with the *preATI* timepoint using Dream, a method designed for repeated-measures designs (58). For each cell subset, we first summed UMI counts across single cells within each sample, yielding a sample-by-gene matrix for that subset. We included only samples with at least 10 cells. We subset the matrix to include samples from *preATI* and a later timepoint of interest and removed lowly expressed genes by retaining only those with counts per million greater than 1 in more than 40% of samples. Because some participants were missing longitudinal samples, either due to the lack of PBMC aliquot or low cell counts in a specific single-cell cluster, differential expression was modeled via Dream (variancePartition package) to accommodate the unbalanced repeated-measures design without data loss. To account for the repeated-measures structure, participant ID was modeled as a random effect, with timepoint included as the primary fixed effect of interest. The CITE-seq experiment was performed in 10 batches, with 3-4 samples from the same timepoint pooled in a batch. To account for the technical noise from the nested batch design, we used the nested-batch term as a random effect (1|timepoint:batch). We transformed the raw counts using the *voomWithDreamWeights* function and built the model using the *dream* function. The significant changes over time were identified using *topTable* function. For a small number of cell subsets, the Dream analysis failed due to too few participants with both timepoints; in these cases, we used DESeq2 (59) to identify genes differentially expressed between pairs of timepoints without accounting for the nested-batch effect.

To identify outcome-specific signals, we compared participants with controller status to those with non-controller status at each timepoint using DESeq2. The same procedure was used for generating a matrix of raw counts as above. We removed lowly expressed genes by retaining only those with counts per million greater than 1 in more than 50% of samples within at least one outcome group, and used the resulting matrix as input to DESeq2, with controller-status as the outcome variable.

To identify differentially active pathways, we used gene-set enrichment analysis (GSEA) (60) for each comparison in the differential expression analysis using Hallmark and Reactome pathway databases. Briefly, we ranked genes by log_2_ fold difference from DREAM or DESeq2, evaluated pathway enrichment using the fgsea R package (v1.18.0), and adjusted the p-values using Benjamini-Hochberg procedure separately for each pathway database. Pathways with adjusted p-values (FDR) < 0.1 were considered significant.

### V(D)J data analysis, quality control, and clonotype expansion analysis

The raw sequencing data for the V(D)J libraries were processed using the Cell Ranger pipeline (v8.0.0; multi command) along with CITE-seq data. Contigs were assembled from reads, aligned, and annotated using the VDJ reference (Ensembl-7.1.0).

For TCR expansion analysis, only the productive alpha and beta chains from the high-quality cells were retained. Chains shorter than 5 or longer than 22 were removed. Because the CDR3 sequences of the beta chain alone accounted for about 98% clonotype diversity, clonotypes were assigned based on beta-chain CDR3 sequences. Only the clonotypes with exactly one beta chain, which accounted for 87% of all captured TCRs, were retained for the downstream analysis. The clonotypes that expanded or contracted over time were identified using Fisher’s exact test for each pair of timepoints of interest within each participant. Although Fisher’s exact test accounts for differences in the total number of observations between samples, it does not account for sampling sparsity arising from incomplete repertoire capture. Because 10x TCR-seq samples only a fraction of the total TCR repertoire, stochastic dropout can inflate false positive rates. To mitigate this effect, the sample with the higher number of TCRs was randomly downsampled to match the size of the other sample prior to Fisher’s exact testing. This downsampling procedure was repeated 100 times with independent random subsampling at each iteration. In each repeat, clonotypes with FDR-adjusted Fisher’s exact test p-values <0.1 and log_2_ fold-changes ≥1 were classified as significantly changed. Clonotypes meeting these criteria in more than 50 of 100 repeats were considered significantly changing clonotypes.

### Intracellular cytokine staining (ICS) and proliferation (CTV)

The magnitude and phenotype of HIV-specific CD4+ and CD8+ T cells was characterized using intracellular cytokine staining and their proliferation was determined using a 6-day *in vitro* peptide stimulation assay using overlapping Clade B HIV peptide pools (NIH HIV Reagent Program, Division of AIDS, NIAID, NIH). These experiments were performed as previously described (13). See **Supp. Table 6** for ICS and CTV panels.

### Mass cytometry (CyTOF)

CyTOF analyses on samples from the *preATI, bNAb, preR*, and *postR* timepoints were performed over the course of five separate experiments as described previously (61). PBMCs were thawed and only samples with >70% viability were used for analysis. 2-4 million cells were stained per panel in two mass cytometry panels (**Supp. Table 6**) following a previously published protocol (62). Manual gating was performed on this dataset as previously described (13).

### Statistical analysis of CyTOF, ICS, and proliferation data

To detect significant changes in immune cell frequencies and phenotypes over time, generalized linear models (GLMs) were fit to the data using the svyglm option from the survey package (v4.4-8). The models were fit with weights set to the total counts and family set to quasibinomial to prevent overdispersion given large weights and low sample size. To test the effect of time on proportions, a Likelihood Ratio Test was performed to compare a full model fit including time and participant-level effects to a reduced model fit with participant-level effects only using the ANOVA function. Similarly, to detect changes in the ICS data over time, GLMs were fit using the glmer function from the lme4 package (v1.1-35.1) using the gaussian family, since weights were not available. Then, an F-test using the Kenward-Roger approximation was performed on a full model fit with time and participant-level effects to a reduced model fit with PID effects only using the KRmodcomp function from the pbkrtest package (v0.5.3). Finally, multiple hypothesis correction was performed separately for each pairwise comparison using the Benjamini-Hochberg procedure. For the proliferation data, the frequency of CTVlo non-naïve CD4+ or CD8+ T cells for samples with available paired datapoints was compared between *preATI* and *preR* using Wilcoxon signed rank testing.

## General

We thank all the UCSF amfAR combination immunotherapy trial participants for making this manuscript and study possible as well the entire SCOPE clinical team. We thank the Gladstone Institutes Genomics Core for library preparation for the CITEseq experiments and the UCSF Center for Advanced Technology for sequencing.

## Author contributions

JAW and RLR conceived of and wrote the manuscript. DAS, RT, SSS, KS, PM, CDG, SI, BJ, MS, MPB, and AND performed experiments and/or generated data for the manuscript. MJP and SGD led and RH, TD, and MCW clinically supported the amfAR trial. JAW, RKP, DAS, GWG, RT, MT, RT, MHS, GKF, PWH, and RLR analyzed data for the manuscript. All authors critically reviewed the manuscript.

## Data and materials availability

CyTOF and sequencing data will be made publicly available. Flow cytometry data will be made available upon request.

