## Supplemental Figures for "Immune activation during broadly neutralizing antibody-mediated HIV suppression prior to post-intervention control"

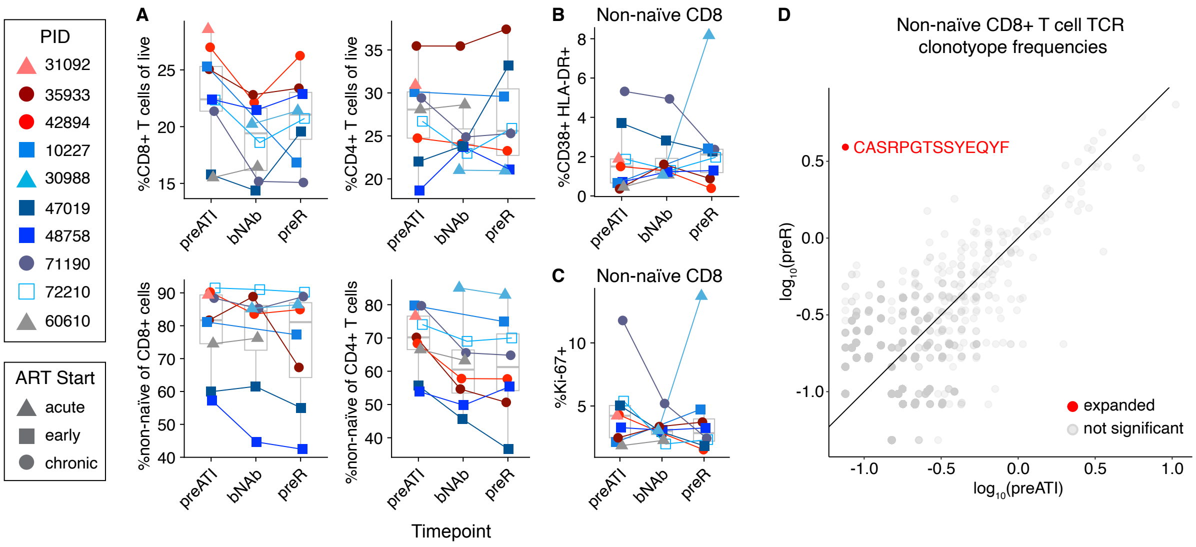


**Supp. Fig. 1: T cell frequency and activation are unchanged under bNAb-mediated HIV suppression.** (A) Frequencies of total and non-naïve CD4+ and CD8+ T cells at the *preATI*, *bNAb*, and *preR* timepoints. (B) Frequency of activated (CD38+HLA-DR+) non-naïve CD8+ T cells at the *preATI*, *bNAb*, and *preR* timepoints. (C) Frequency of proliferative (Ki-67+) non-naïve CD8+ T cells at the *preATI*, *bNAb*, and *preR* timepoints. (D) An example plot showing non-naïve CD8+ T cell TCR clonotype frequencies from 10x scRNA/TCRseq data at the *preR* (y-axis) versus *preATI* (x-axis) timepoints (1 clonotype significantly expanded in PID 30988 at *preR* relative to *preATI*). For A-C: data were compared using linear mixed effects models with Benjamini-Hochberg correction; no comparisons were statistically significant ($p_{adj}<0.05$). For D: significant expansion was defined as $p_{adj}$ <0.1 and $\log_2$ fold-change ≥1 in more than 50 of 100 repeats. Box plots indicate median (line) and interquartile range (boxes).

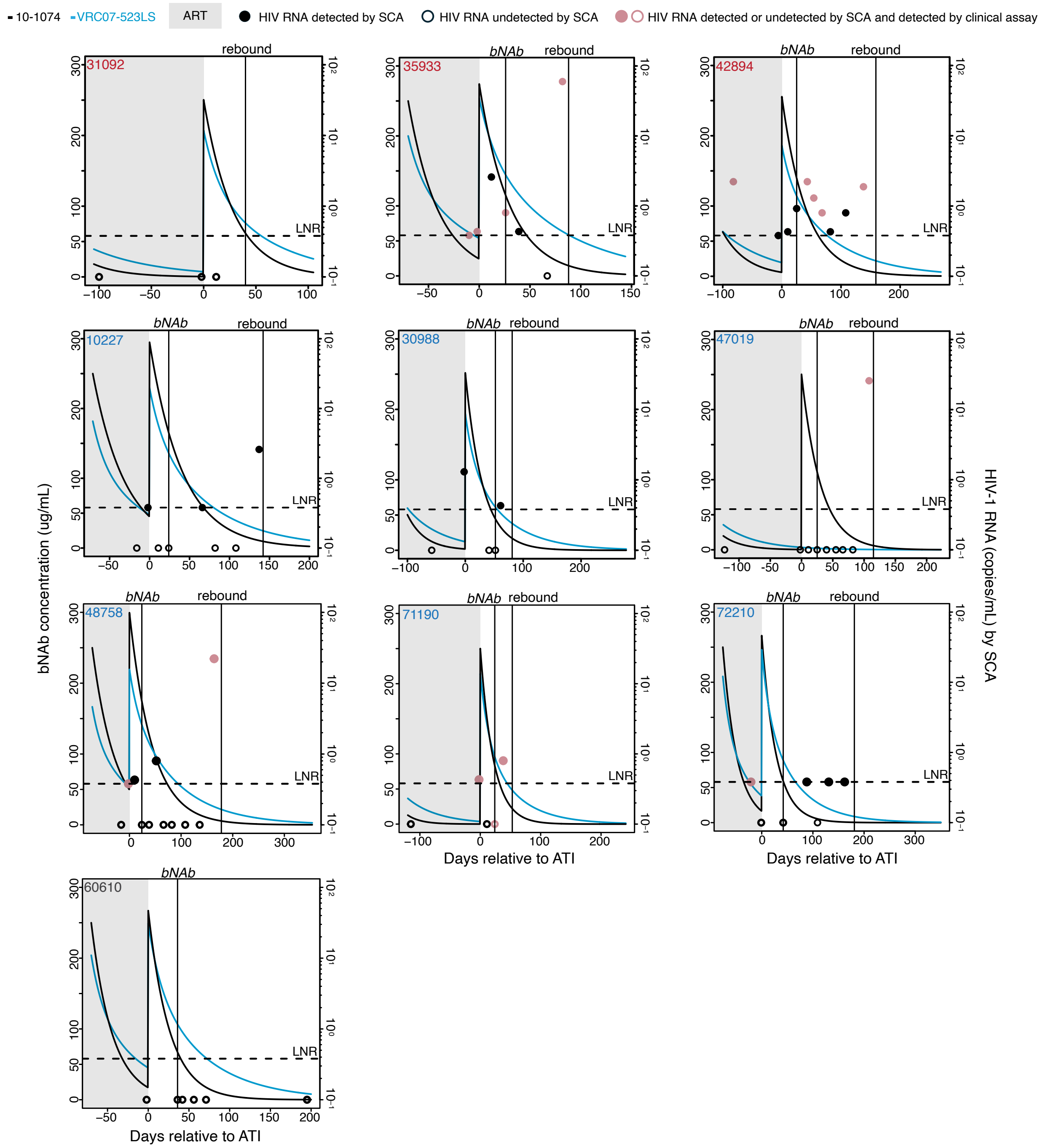


**Supp. Fig. 2: Minimal residual plasma viremia detectable during bNAb-mediated HIV suppression.** bNAb levels (10-1074 [black] and VRC07-523LS [blue]) and HIV-1 RNA levels (circles) by single copy assay for each participant at the indicated day relative to ATI (day 0). The *bNAb* timepoint and time of rebound are indicated with a solid vertical line for each participant. For the viral loads, black indicates HIV-1 RNA was not detected by clinical assay at that timepoint and rose indicates HIV-1 RNA was detected by clinical assay. Solid circles indicate HIV-1 RNA was detected by single copy assay at that timepoint versus empty circles indicate HIV-1 RNA was not detected by single copy assay. A shaded grey background indicates ART suppression.

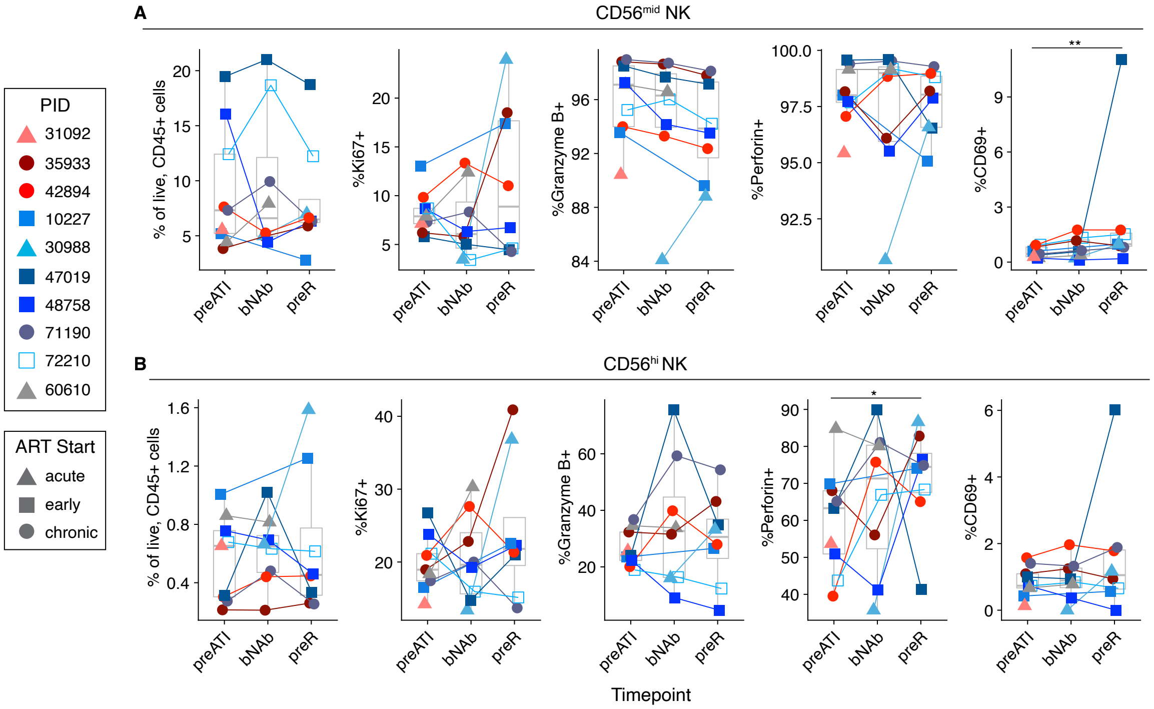


**Supp. Fig. 3: Additional innate immune activation markers.** Ki-67+, Granzyme B (GzB)+, Perforin+, CD25+, and CD69+ of (A) CD56$^{mid}$ or (B) CD56$^{hi}$ NK cells at the *preATI*, *bNAb*, and *preR* timepoints. Data were compared using linear mixed effects models with Benjamini-Hochberg correction. * = $p_{adj}$<0.05, ** = $p_{adj}$< 0.01. Box plots indicate median (line) and interquartile range (boxes).

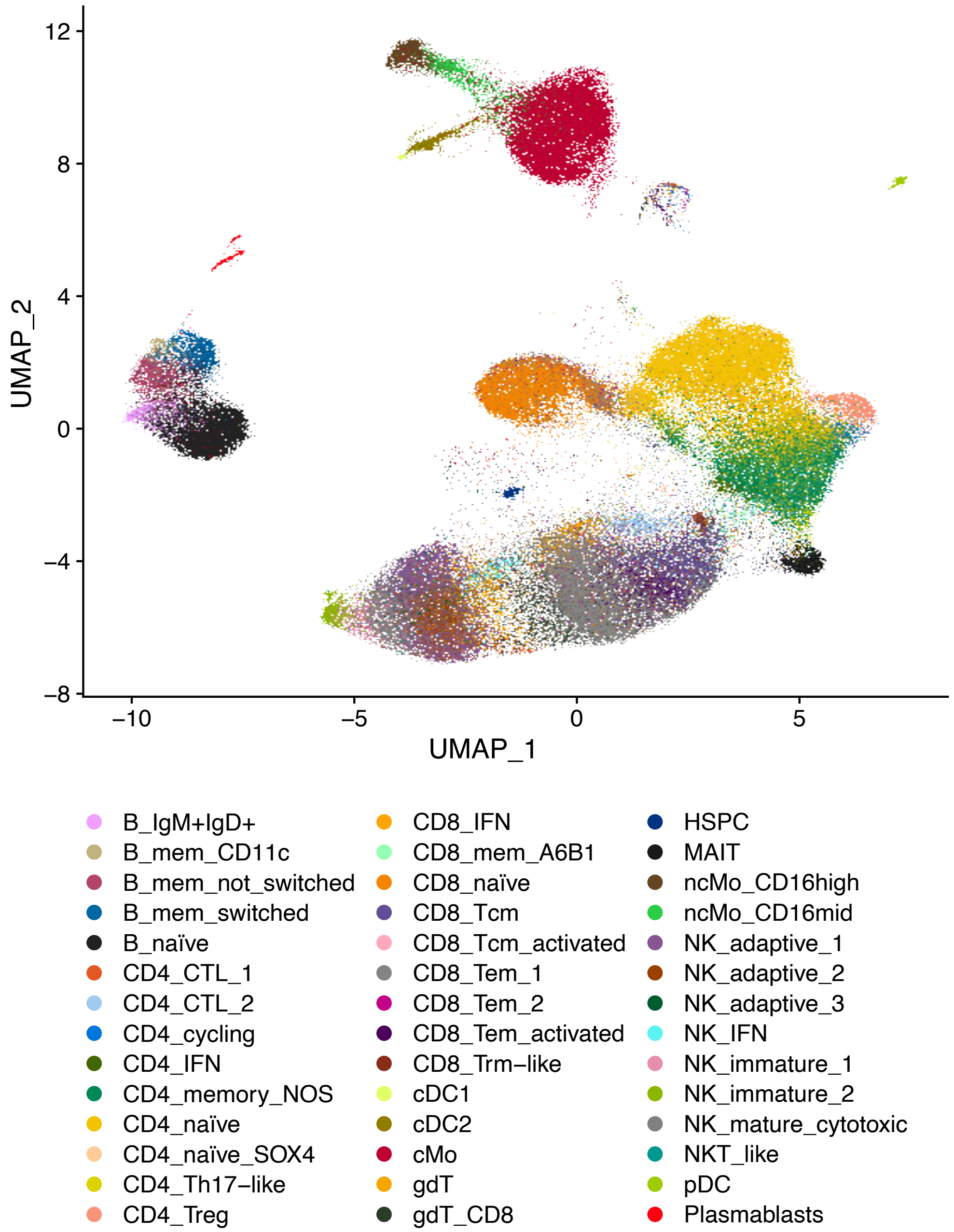


**Supp. Fig. 4: UMAP of all 42 cell type subclusters identified using CITEseq.**

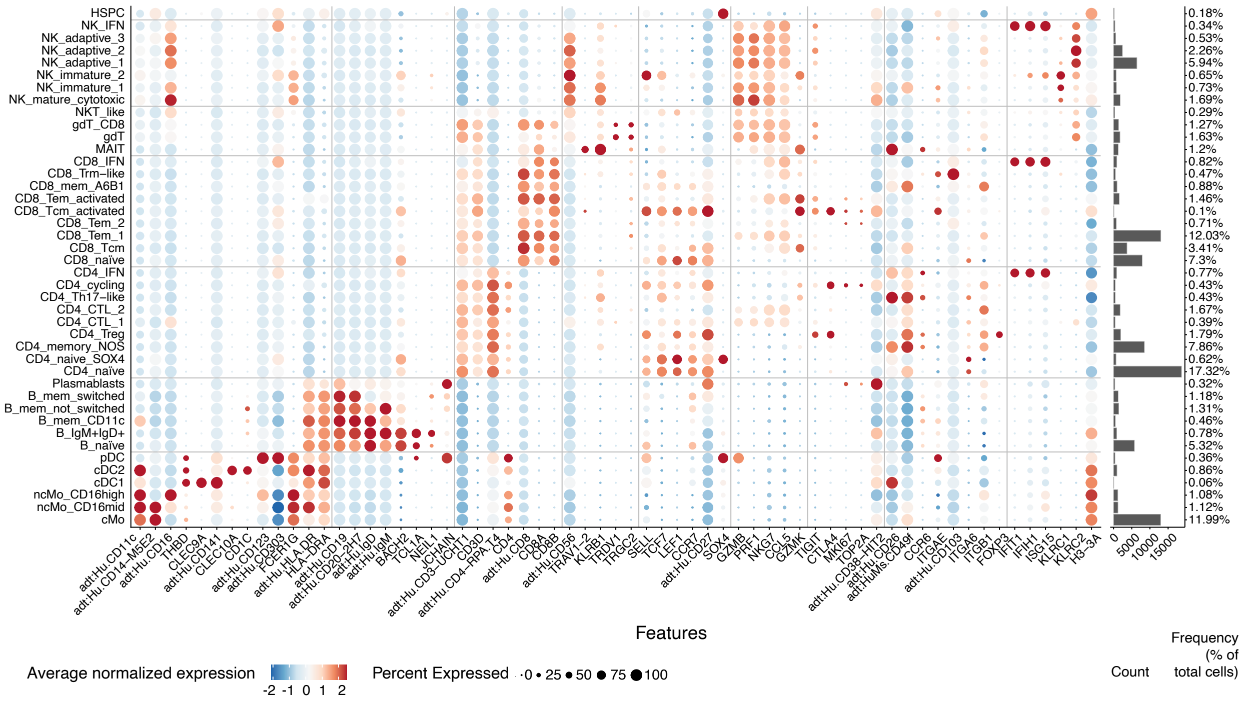

**Supp. Fig. 5: Gene transcript and protein expression data used to identify CITEseq cell type clusters.** Abbreviations: hematopoietic stem cell (HSPC), natural kill cell (NK), interferon (IFN), γδ T cell (gdT), mucosal-associated invariant T cells (MAIT), T resident memory (Trm), memory (mem), effector memory (em), central memory (cm), cytotoxic lymphocyte (CTL), not otherwise specified (NOS), plasmacytoid dendritic cell (pDC), conventional dendritic cell (cDC), non-classical monocyte (ncMO), classical monocyte (cMo).

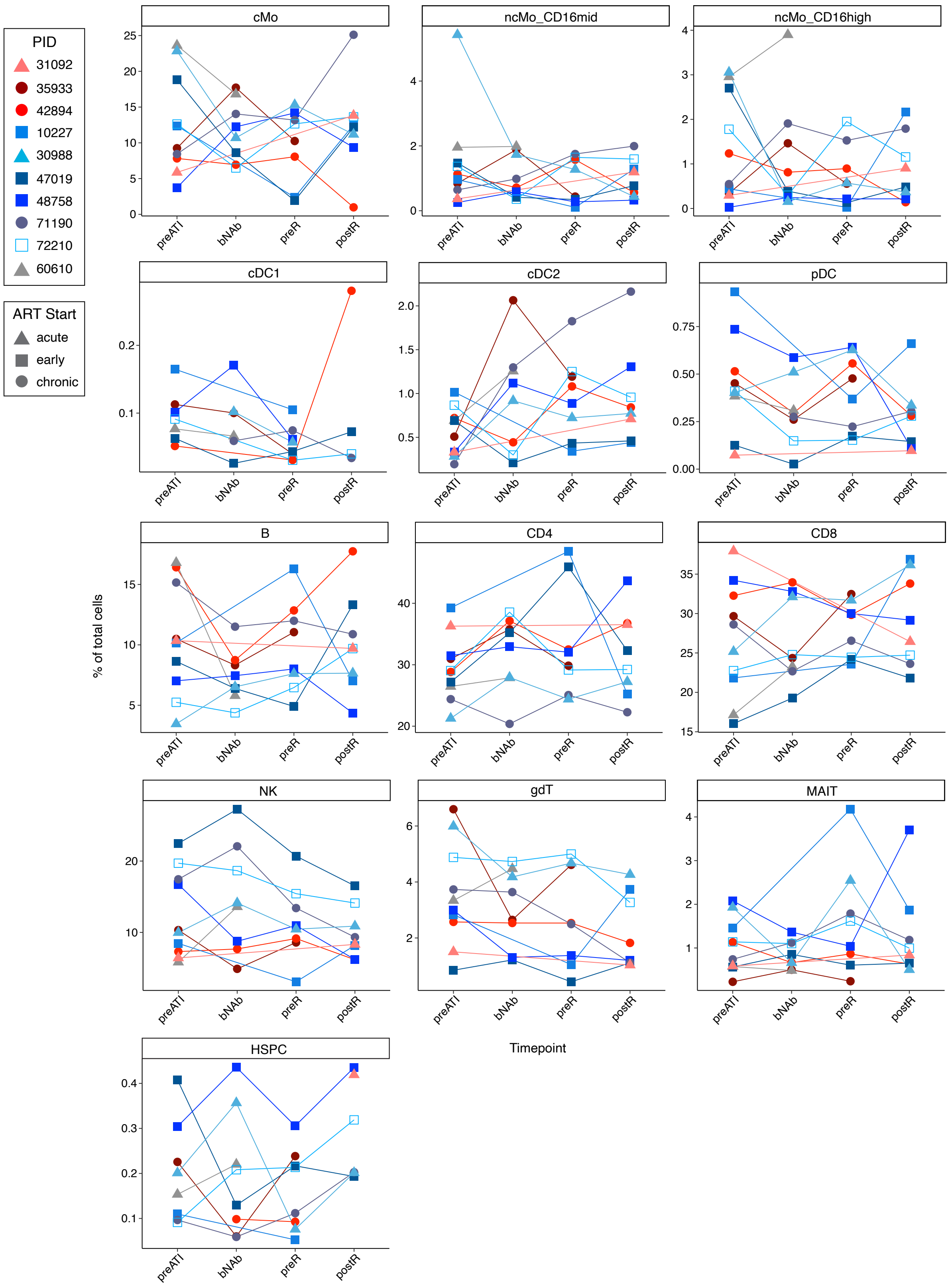


**Supp. Fig. 6: Frequencies of landmark populations under bNAb-mediated HIV suppression by CITEseq.** Frequencies are of total cells.

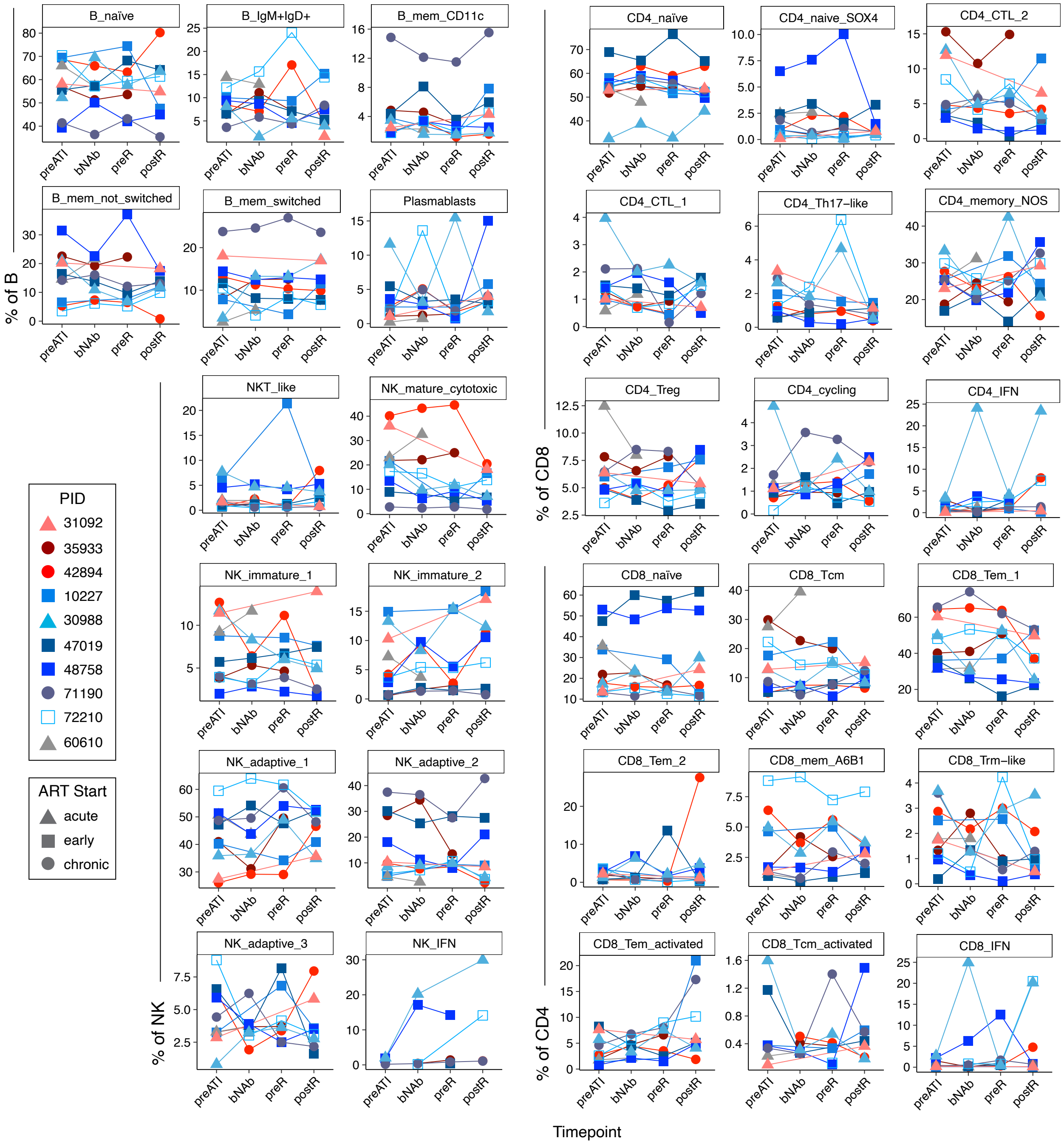


**Supp. Fig. 7: Frequencies of cell type subclusters under bNAb-mediated HIV suppression by CITEseq.** Frequencies are of parent population (B cells, NK cells, CD4+ T cells, CD8+ T cells).

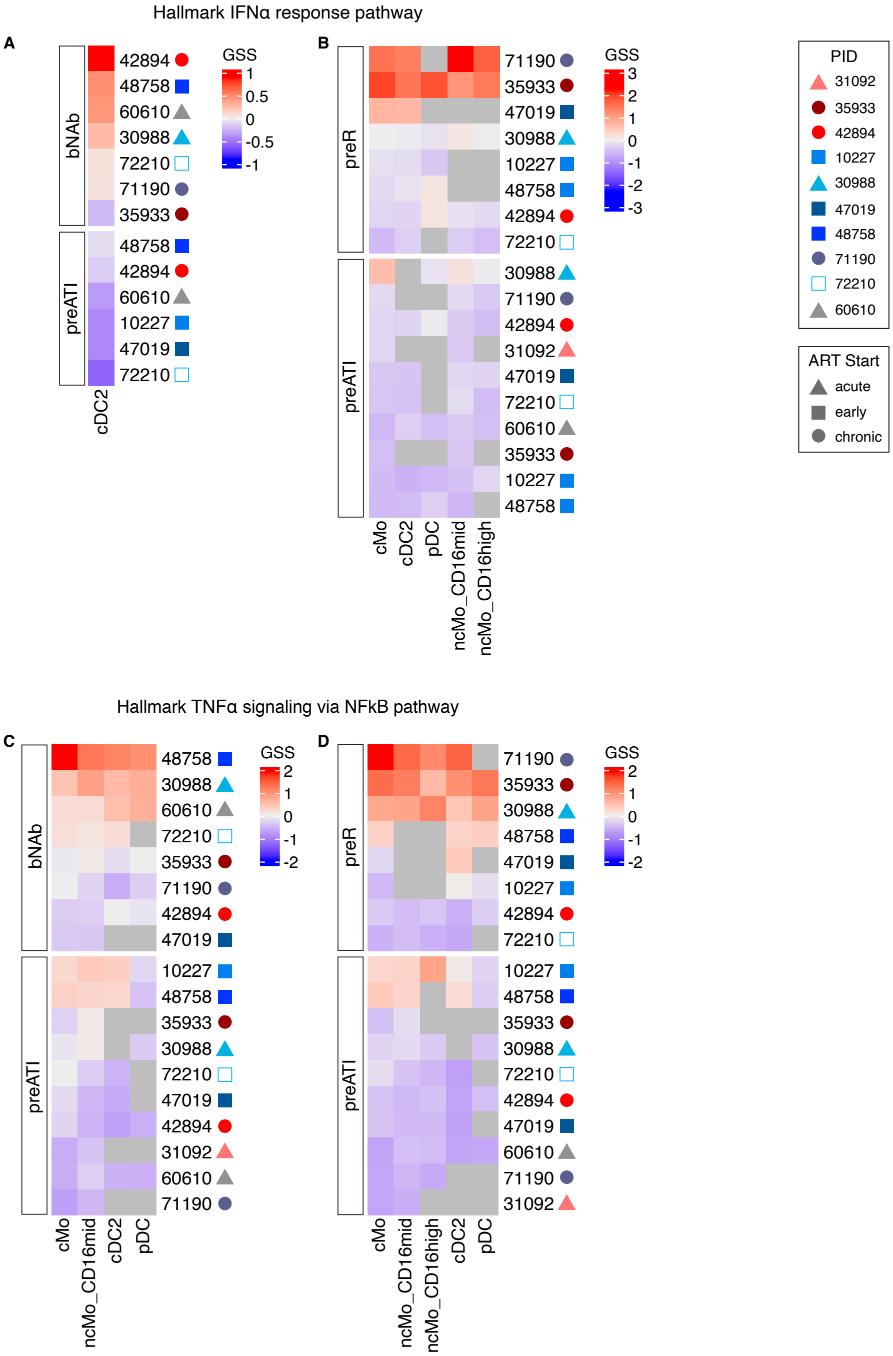


**Supp. Fig. 8: IFNα and TNFα signaling gene signature scores by participant for between timepoint comparisons.** Gene signature scores (GSS) for the Hallmark Interferon (IFN)α response and Hallmark TNFα signaling via NFκB pathways for each indicated cell type by participant at the (A, C) *bNAb* and *preATI* and (B, D) *preR* and *preATI* timepoints. Gene signature score reflects the mean of the Z-scored expression of leading-edge genes of the relevant pathway in the indicated sample/cell type. Grey boxes indicate no data.

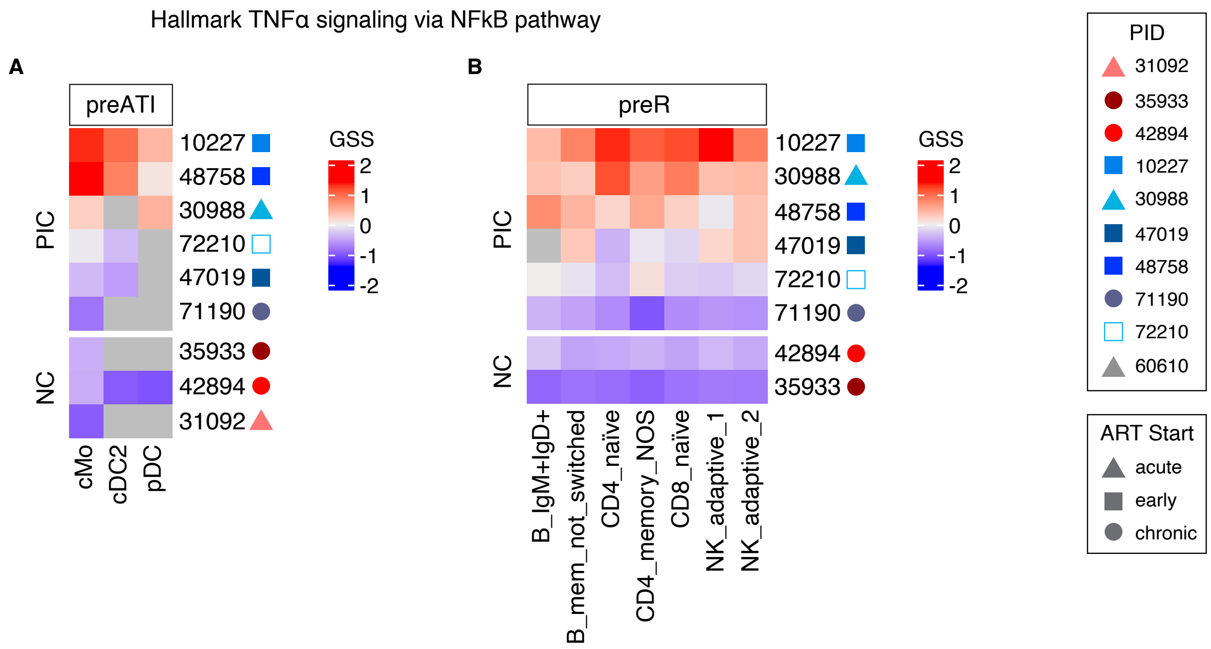


**Supp. Fig. 9: TNFα signaling gene signature scores by participant for controller versus non-controller comparisons.** Gene signature scores (GSS) for each indicated cell type by participant at the (A) *preATI* and (B) *preR* timepoints. Gene signature score reflects the mean of the Z-scored expression of leading-edge genes of the Hallmark TNFα signaling via NFκB pathway in the indicated sample/cell type. Grey boxes indicate no data. NC = non-controller, PIC = post-intervention controller.
